# md_harmonize: a Python package for atom-level harmonization of public metabolic databases

**DOI:** 10.1101/2022.12.08.519680

**Authors:** Huan Jin, Hunter N.B. Moseley

## Abstract

**Summary:** A big challenge to integrating public metabolic resources is the use of different nomenclatures by individual databases. This paper presents md_harmonize, an open-source Python package for harmonizing compounds and metabolic reactions across various metabolic databases. md_harmonize utilizes a neighborhood-specific graph coloring method for generating a unique identifier for each compound via atom identifiers based on the compound structure. The resulting harmonized compounds and reactions can be used to construct metabolic networks and models for various downstream analyses, including metabolic flux analysis.

**Availability:** The md_harmonize package is implemented in Python and freely available at https://github.com/MoseleyBioinformaticsLab/md_harmonize

**Supplementary information:** Supplementary data are available at https://doi.org/10.6084/m9.figshare.21699683.

## 1. Introduction

Metabolic reprogramming has been recognized as a phenotypic hallmark of cancer (Faubert, et al., 2020). Large-scale characterization of metabolic phenotypes of cancers can help clarify biological mechanisms of metabolic diseases and develop novel and effective therapeutics (DeBerardinis and Chandel, 2016). Stable isotope tracing is an essential tool in deciphering metabolic mechanisms by demultiplexing metabolic fluxes (You, et al., 2014). With the rapid development of analytical methodologies, especially nuclear magnetic resonance spectroscopy (NMR) and mass spectrometry (MS) (Fan, et al., 2012), large volumes of high-quality isotopic labeled metabolic profiles are being generated. To derive meaningful biological interpretations from the metabolic datasets, a necessary first step is to construct reliable metabolic models (Jin and Moseley, 2019) via a comprehensive atom-resolved metabolic network, which requires harmonization of compounds and atom-resolved reactions from public metabolic resources (Altman, et al., 2013). Metabolic databases, like KEGG (Kyoto Encyclopedia of Genes and Genomes) and MetaCyc, contain either atom transformation patterns between reactant-product pairs (Kotera, et al., 2004) or direct atom mappings for reactions (Latendresse, et al., 2012), which greatly contribute to the construction of an atom-resolved metabolic network. However, the lack of a uniform identity, especially for the atom identifiers, is a big challenge in integrating metabolic databases (Jin, et al., 2020; Jin and Moseley, 2021; Poolman, et al., 2006; Powers, 2009).

A neighborhood-specific graph coloring method was developed to generate unique identifiers for every atom and the corresponding compound based on the structure (Jin, et al., 2020). This method requires only the molfile representation of the compound, which is available in most metabolic databases. These unified compound identifiers help facilitate both compound and reaction harmonization across metabolic databases.

Here, we developed a Python package md_harmonize, utilizing the neighborhood-specific graph coloring method, to facilitate harmonization of compounds and atom-resolved metabolic reactions across various metabolic databases. This package integrates data standardization, aromatic substructure detection, and compound and reaction harmonization together. It provides a user-friendly command-line interface for easy implementation.

## 2. md_harmonize package

### 1. md_harmonize package overview

As shown in Fig.1, the md_harmonize Python package is composed of several modules. The compound.py module defines the basic elements for compound construction, composed of ‘Atom’, ‘Bond’, and ‘Compound’ classes. The aromatics.py module contains ‘AromaticManager’ class that is responsible for aromatic substructure construction and detection. The algorithm Biochemically Aware Substructure Search (BASS) (Mitchell, et al., 2014) for aromatic substructure detection is implemented in the ‘BASS.pyx’ module. Data structure for constructing reaction is documented in the ‘reaction.py’ module. The ‘harmonization.py’ module takes charge of harmonizing compounds as well as reactions. The harmonized results are represented by ‘HarmonizedCompoundEdge’ and ‘HarmonizedReactionEdge’ objects. The ‘__main__.py’ module provides the command-line interface to perform data standardization, aromatic substructure detection, and compound and reaction harmonization, which is implemented with the ‘docopt’ Python library. The other modules ‘tools.py’, ‘KEGG_database_scraper.py’, ‘KEGG_parser.py’, ‘Meta-Cyc_parser.py’, ‘openbabel_utils.py’, and ‘supplement.py’ define auxiliary tools for reading and writing files, scraping and parsing data, standardizing molfile representation via Open Babel (O’Boyle, et al., 2011), and processing variables.

**Fig. 1.**
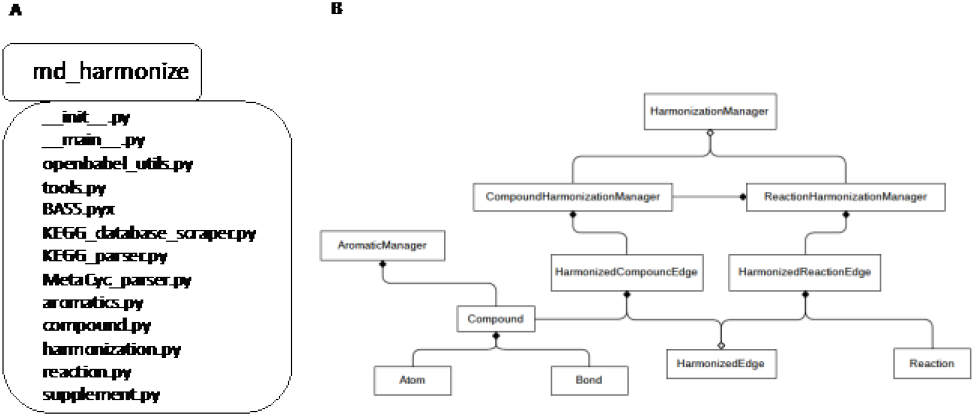
Organization of md_harmonize package presented with UML diagrams: A) UML package diagram of the md_harmonize Python library; B) UML class diagram of the md_harmonize Python package.

### 2. md_harmonize package interface

The md_harmonize package provides a simple command-line interface to perform data standardization, aromatic substructure detection, and compound and reaction harmonization. Fig. 2 shows version 1.0 of the command-line interface.

**Fig. 2.**
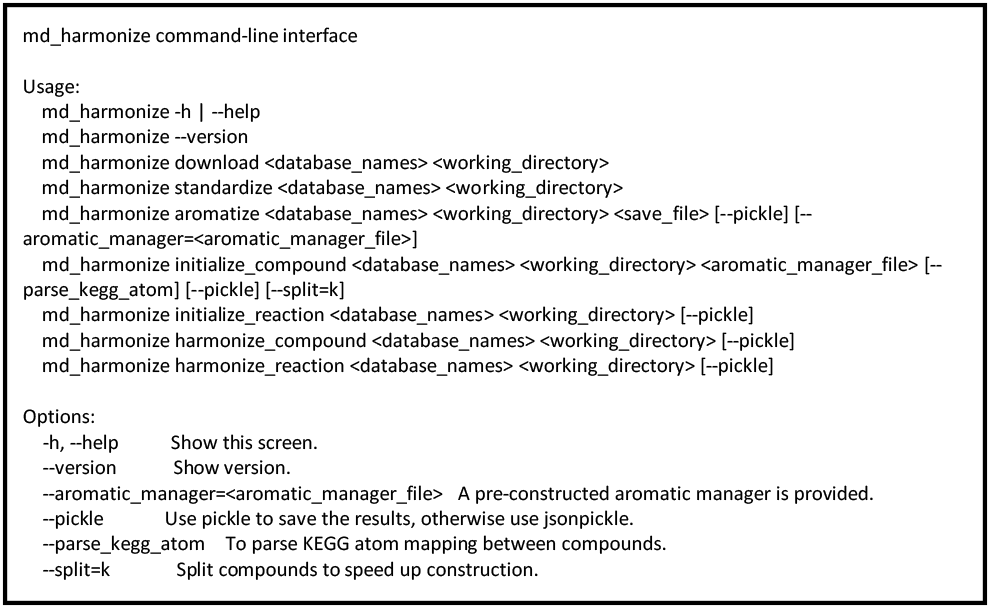
Command line interface of md_harmonize package.

## 3. Application

In our prior work, we demonstrated compound and reaction harmonization of KEGG and MetaCyc using a predecessor prototype of md_harmonize (Jin and Moseley, 2021). Here, we used three major metabolic databases, KEGG, MetaCyc, and Human Metabolome Database (HMDB) to evaluate the results of compound harmonization generated by md_harmonize. Some compounds in HMDB have direct KEGG and/or MetaCyc references. We extracted all the HMDB-KEGG and HMDB-MetaCyc compound pairs from HMDB, then tested if those pairs can be detected by md_harmonize. The results are shown in Table 1. Based on the direct references of HMDB, 6814 KEGG-HMDB compound pairs and 2652 MetaCyc-HMDB compound pairs were detected. With the md_harmonize, we discovered 8644 KEGG-HMDB compound pairs and 7271 MetaCyc-HMDB compound pairs. About 5358 KEGG-HMDB compound pairs and 1868 MetaCyc-HMDB compound pairs can be cross-validated, indicating that md_harmonize was able to detect thousands more compound pairs.

**Table 1.** Comparison of compound harmonization.

| Databases | HMDB extracted | md_harmonize detected | Overlap |
| --- | --- | --- | --- |
| KEGG | 6814 | 8644 | 5358 |
| MetaCyc | 2652 | 7271 | 1868 |

On the other hand, the md_harmonize method missed about 1456 KEGG-HMBD compound pairs and 784 MetaCyc-HMDB compound pairs indicated by HMDB references. We further investigated the causes of missed detection (Table 2). About 232 KEGG compound references and 111 MetaCyc compound references are invalid, either no molfile representation or incorrect compound identifier provided. We then compared the compound molecular formulas, and found that 793 KEGG-HMDB compound pairs and 557 MetaCyc-HMDB compound pairs have inconsistent molecular formulas.

**Table 2.** Categorization of missed compound pairs by md_harmonize.

| Category | KEGG | MetaCyc |
| --- | --- | --- |
| Invalid references | 232 (15.93%) | 111 (14.15%) |
| Inconsistent formulas | 793 (54.46%) | 557 (71.05%) |
| Other | 431 (29.61%) | 116 (14.80%) |
| Total | 1456 | 784 |

The remaining 431 KEGG-HMDB compound pairs and 116 Meta-Cyc-HMDB compound pairs are likely caused by inconsistent compound structures as illustrated in Fig 3. Also, circular-linear interchangeable structures would not be detected, since md_harmonize requires reaction descriptions from both databases for reliably detecting such pairs and HMDB does not have the required reaction descriptions.

**Fig. 3.**
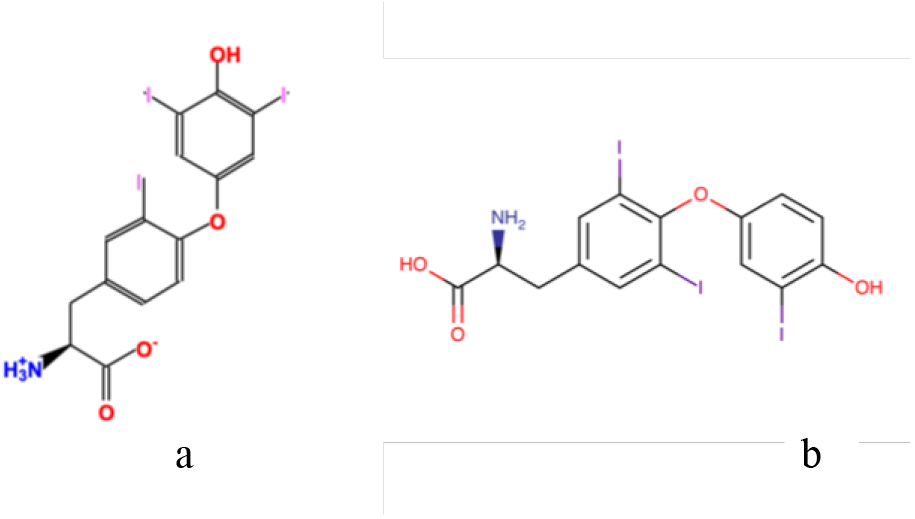
Example of incorrect compound pair indicated by HMDB reference with different structure representations. a) MetaCyc CPD-10813; b) HMDB HMDB0000265.

## 4. Conclusions

The md_harmonize Python package provides data structures and algorithms for harmonizing public metabolic databases. As demonstrated by the added HMDB functionality of md_harmonize, the modular design, common data structures, and base parsing functionality enables highly flexible extensions to be added to support additional metabolic and compound databases. Additionally, the harmonized structured results generated by md_harmonize can be easily used to construct integrated metabolic networks and associated atom-resolved metabolic models.

